# Altered excitability of dI3 neurons regulates hindlimb motor tone and locomotor recovery after spinal cord injury

**DOI:** 10.1101/2025.09.03.674123

**Authors:** Alex M. Laliberte, Shahriar Nasiri, Matthew Redmond, Sara Goltash, Riham Khodr, Nikki Lalonde, Tuan V Bui

## Abstract

Recovery of motor function after spinal cord injury is limited in mammals. Reactivation of locomotor circuits does occur, but primarily through the activation of sensorimotor pathways in the context of locomotor training. Previous investigations have shown that dI3 neurons, a developmentally-defined population of pre-motor, glutamatergic interneurons, are indispensable for this process. However, it remains unclear how dI3 neurons are recruited during locomotor recovery, and whether they could be leveraged to improve locomotor function following spinal cord injury. Herein, we investigated how the excitability of dI3 neurons influences locomotor behaviour and recovery after spinal cord injury. In T9-T10 transected mice, we found that acute chemogenetic silencing of dI3 neurons leads to immediate loss of hindlimb motor tone, and significant reduction in stepping during treadmill locomotion. Conversely, regular chemogenetic stimulation of dI3 neurons led to transient increases in hindlimb motor tone early after injury, but ultimately reduced hindlimb motor tone and locomotor recovery over the long term. These chronic changes resulting from dI3 neuron stimulation were associated with the absence of expression of the constitutive 5-HT2C-R isoform, potentially representing a homeostatic mechanism for the regulation of dI3 excitability following spinal cord injury. Given these findings, we hypothesized that dI3 stimulation’s effects on motor tone, while insufficient to drive locomotor function alone, may promote stepping improvements when locomotor rhythm-generating circuits are active. The addition of quipazine, a serotonergic agonist known to facilitate locomotor rhythmogenesis, in combination with dI3 stimulation, significantly improved locomotor function, while also mitigating the long-term reduction in treadmill stepping associated with dI3 stimulation alone. In aggregate, our results suggest that hyper-excitable dI3 neurons are involved in the maintenance of motor tone after spinal cord injury, possibly through a 5-HT2C-R-dependent mechanism, and further show that the selective stimulation of dI3 neurons could enhance the recovery of locomotor function following spinal cord injury.

## INTRODUCTION

Recovery of lost motor function after spinal cord injury is limited. Reactivation of spinal locomotor circuits that have been disconnected from their descending inputs is possible, particularly through sensory inputs (Barbeau and Rossignol, 1987; Edgerton and Roy, 2009; Sławińska et al., 2012). This engagement of spinal locomotor circuits after spinal cord injury is evidenced by locomotor activity of spinalized animals when positioned on a treadmill. This locomotor activity is purported to be generated by the activation of sensory inputs related to proprioception and low-threshold mechanoreception that feed back into central spinal locomotor circuits and produce cyclic leg movements with similar rhythms and patterns of coordination that are the hallmarks of locomotor activity (Bouyer and Rossignol, 2003; Sławińska et al., 2012; Takeoka et al., 2014; Takeoka and Arber, 2019).

The capacity of sensory inputs to activate spinal locomotor circuits varies amongst species. Spinalized mammals such as cats, dogs, rats and mice, which generally experience moderate recovery of locomotor function after SCI, still require several weeks of recovery post-injury before demonstrating weight support and locomotor activity on a treadmill (Barbeau and Rossignol, 1987; Sławińska et al., 2012; Santos et al., 2024). While it is expected that plasticity plays a central role in this process, the precise cellular substrates and molecular mechanisms involved are not well understood.

In mice, spinal dI3 neurons have been implicated with the recovery of locomotor function after spinal cord injury. This population of spinal neurons arises from the dp3 progenitor domain and is defined, in part, by Isl1 expression. dI3 neurons reside mainly in the deep dorsal horn and intermediate laminae at the interface between sensory and motor circuits within the spinal cord (Bui et al., 2013). Notably, these spinal interneurons integrate proprioceptive and mechanoreceptive information and they project ipsilateral glutamatergic inputs to motoneurons (Bui et al., 2013) and spinal locomotor circuits (Bui et al., 2016). Genetic silencing of VGLUT2 in Isl1-expressing cells, which include dI3 neurons, substantially impairs the recovery of locomotor function in spinalized mice (Bui et al., 2016). Conversely, dI3 neuron silencing has only subtle effects on intact mice during treadmill locomotion (Bui et al., 2016), suggesting that the contribution of dI3 neurons to locomotion may be context-dependent. As such, it remains unclear how dI3 neurons are recruited during locomotor recovery, and whether they could be leveraged to improve locomotor function following spinal cord injury.

This study employs optogenetic and chemogenetic approaches to investigate how dI3 neurons contribute to the recovery of locomotor function in spinalized mice. Using selective activation or inhibition of dI3 neurons in spinalized transgenic mice, along with electrophysiological experiments, we demonstrate that hyperexcitable dI3 neurons are crucial for maintaining motor tone and facilitating the activation of locomotor circuits following injury. Surprisingly, chronic chemogenetic excitation of dI3 neurons impaired the late-stage recovery of locomotor function observed in control spinalized mice, potentially indicating that chemogenetic dI3 stimulation leads to reduced excitability through homeostatic mechanisms. Immunostaining data highlight the expression of the 5-HT2C receptor in dI3 neurons, a receptor known to adopt a constitutively-active isoform following spinal cord injury (Murray et al., 2010), as a potential mediator of this effect. Notably, co-administration of quipazine, a serotonergic agonist, alongside chemogenetic excitation of dI3 neurons, enhanced locomotor performance and mitigated some of the long-term negative effects on recovery.

Together, these findings demonstrate that dI3 neuron excitability plays a key role in regulating motor tone after spinal cord injury. Because motor tone below the injury level can be either beneficial or detrimental depending on its degree, we propose that precise modulation of dI3 neuron excitability may be essential for optimizing locomotor recovery while minimizing the risk of hypertonia.

## MATERIALS AND METHODS

### Animals

All animal procedures were approved by the University of Ottawa Animal Care Committee and conform to the guidelines put forth by the Canadian Council for Animal Care. In order to target gene expression to dI3 neurons, we generated a hybrid transgenic mouse line based on the combination of Isl1-Cre (RRID:IMSR_JAX:024242) and Vglut2-Flp (RRID:IMSR_JAX:030212) mouse lines. This dual-recombinase driver line was then bred to the corresponding dual-recombinase-dependent reporter lines. For the expression of the inhibitory DREADD receptor - hM4Di - dI3 driver mice were bred to RC::FPDi dual-recombinase responsive fluorescent/DREADD mice (RRID:IMSR_JAX:029040), alternatively noted as Cre^ON^-Flp^ON^-hM4Di herein. For optogenetic experiments, dI3 driver mice were bred to the Ai80(RCFL-CatCh)-D (RRID:IMSR_JAX:025109) transgenic line of mice. For simplicity, these mice will be referred as dI3^CatCh^ herein. These mice conditionally express the blue light-sensitive CatCh channels only in cells that are Isl1+ and Vglut2+. This conditional expression limits the expression of CatCh to a population of spinal dI3 neurons (Bui et al., 2016) and a subset of sensory neurons involved in mechanical nociception and thermoception (Lagerstrom et al., 2010; Goltash et al., 2025). To express the fluorescent reporter tdTomato, dI3 driver mice were similarly bred to the Ai65(RCFL-tdT) (RRID:IMSR_JAX:021875) line.

### Intraspinal AAV microinjection

Bilateral intraspinal injections were performed to deliver adeno-associated virus (AAV) vectors into the lumbar spinal cord. Six-week-old mice were anesthetized using isoflurane (4% induction, 1–2% maintenance in oxygen) and positioned in a stereotaxic frame. A midline dorsal incision was made over the thoracolumbar vertebrae, and muscle tissue was carefully retracted to expose the vertebral column. Interlaminar windows were identified between the T11–T12, T12–T13, and T13–L1 vertebrae, corresponding to the approximate location of the lumbar spinal cord.

A quartz micropipette (pulled to a ∼30–50 µm tip diameter) was attached to a removable-needle Hamilton syringe using a compression fitting, and backfilled with the paraffin oil. The vector solution was drawn into the syringe by placing the needle tip into a drop of the vector solution and withdrawing at a rate of 1 μL/min using a Harvard Apparatus Pump 11 Elite Nanomite mounted on a micromanipulator. Using the same system, 300 nL of AAV was injected per site at a depth of ∼650-700 µm from the dorsal surface of the spinal cord at a rate of 100 nL/min. Injections were made bilaterally at each interlaminar window, for a total of six injection sites per animal. The micropipette was left in place for 1–2 minutes post-injection to minimize backflow before being slowly withdrawn. Muscle and skin were sutured in layers, and animals were monitored postoperatively until full recovery. Analgesics (buprenorphine, 0.1 mg/kg, s.c.) were administered pre- and post-surgery according to institutional animal care protocols.

### Surgical details of spinal cord injury

Complete transections of the spinal cords were performed at T9-T10 under isoflurane anesthesia. This injury interrupts all descending supraspinal pathways. Sterile surgical foam (Surgifoam, Johnson & Johnson Medtech) was placed in the spinal space to reduce bleeding and to prevent any re-growth of the axons across the lesion. Animals were individually housed and given analgesic (Buprenorphine, Ceva Animal Health, 0.1 mg/kg S.C.) for three days. Following a one-week recovery period, mice were anesthetized with isoflurane and custom-built EMG electrodes were implanted based on a protocol by Turgay Akay (Akay et al., 2006; Bui et al., 2013). After shaving the neck and hindlimbs of the mice, small incisions were made into the skin at the neck area and at the hindlimbs. The electrodes were led under the skin from the neck to the leg incisions using needles and implanted into the left and right tibialis anterior and gastrocnemius/soleus muscles. The incisions on the hindlimbs were closed with 7-0 prolene sutures, and the connector was stitched to the skin near the neck incision using 5-0 silk sutures (Ethicon) and mice were given analgesic and allowed to recover for 3 days.

### Electrode fabrication

The electrodes were made using multi-stranded, Teflon coated annealed stainless-steel wire (#793200, A-M systems). Four pairs of electrodes were prepared for each mouse. For each electrode pair, two pieces of wires were cut about 15 cm in length and twisted together to make a knot. On the first strand, 1mm distal to the knot, a 1mm section of the Teflon coating was stripped off. On the second strand, 3mm distal to the knot, 1 mm of the insulation was stripped off so that the bare regions from each wire were separated by about 2 mm. Next, about 8-9 mm of coating was removed from one end of the wires, dipped in glue to prevent the exposed wires from fraying and crimpled into the shaft of a 27-gauge needle. The opposite ends of the wires were cut about 8 cm from the knot, stripped of their insulation, and soldered to an eight-pin miniature connector. The underside of the connector was covered with epoxy (Devcon 5 min Epoxy Gel) to electrically insulate the wire insertions that were made into the connector and the epoxy was extended about 2 mm beyond the edges of the connector. Small holes were drilled into the edges of the epoxy to provide anchors for suturing the connector into the skin.

### Treadmill Training and Assessment

To assess locomotor recovery following spinal cord injury (SCI), mice underwent training sessions were conducted three times per week, with each session lasting 15 minutes. To acclimate the animals to treadmill walking, training began two weeks prior to SCI surgery. During this acclimation period, mice were placed on the treadmill and encouraged to walk at a constant speed appropriate for their ability (typically 8–15cm/s). Following SCI, mice were given a 1-week recovery period without treadmill exposure. Treadmill training then resumed and continued three times weekly for the remainder of the experimental timeline. During post-injury sessions, treadmill speed was reduced to ∼ 7cm/s, and partial body weight support was provided by the experimenter to facilitate stepping movements and encourage participation. All training was conducted during the light phase of the light/dark cycle, and animals were monitored for signs of fatigue or distress. On weeks 2, 4, 6, and 8 post-injury, an additional treadmill session was provided to assess locomotor function. These sessions included both weight-assisted and unassisted treadmill tests to assess stepping and motor tone/weight bearing, respectively. Sessions were recorded using a Basler acA640-750um camera at a frame rate of 60 fps.

### Pose Estimation and Kinematic Analysis

Hindlimb kinematics were quantified using DeepLabCut v3.0.0rc7 (Mathis et al., 2018; Nath et al., 2019) a deep learning-based markerless pose estimation tool. Anatomical landmarks on the visible hindlimb – including the iliac crest, hip, knee, ankle, metatarsophalangeal (MTP) joint, and toe – were manually labeled across a representative subset of video frames. A neural network using the ResNet-50 architecture was trained for 200 epochs, and the best-performing model snapshot (based on validation error) was selected for further refinement. Outliers in predicted labels were identified using DeepLabCut’s uncertainty algorithm, and a single round of refinement was performed. The network was then retrained for an additional 100 epochs, with the updated best-performing snapshot selected for final analysis. A trained model was applied to a full video dataset for a given experiment, and ARIMA filtering was used to smooth tracking jitter and improve overall temporal consistency. The resulting joint coordinate data were processed using custom Python scripts to calculate joint angles and quantify stepping.

### *In Vivo* Chemogenetic Modulation of dI3 Activity

To modulate neuronal activity in vivo, mice received targeted expression of Designer Receptors Exclusively Activated by Designer Drugs (DREADDs). Either hM4Di (inhibitory) or hM3Dq (excitatory) DREADDs were expressed via transgenic breeding or delivered via adeno-associated virus (AAV) to target the dI3 neuronal population as described above. To activate DREADDs, the selective agonist JHU37160 dihydrochloride (Hello Bio) was dissolved in sterile saline and administered via subcutaneous injection at a dose of 0.5 mg/kg, 15 minutes prior to the start of behavioral testing. To allow full drug washout, a minimum interval of 48 hours was maintained between treatment sessions. Mice underwent repeated testing under different treatment conditions, which included JHU37160, saline vehicle control, and quipazine (where applicable). Treatment order was randomized across animals to control for potential sequence effects. To prevent experimenter bias, treatments were prepared and coded by a third party, and the experimenter conducting behavioral assessments was blinded to treatment identity throughout data collection.

### Ex-vivo electrophysiology

Extracellular ventral and dorsal root recordings via suction electrodes were amplified 1000X in differential mode, bandpass-filtered (1 Hz to 1 KHz) using a differential amplifier (AM Systems model 1700), and acquired at 10 kHz (Digidata 1550B, pClamp 10 software, Molecular Devices). Following anaesthesia with sodium pentobarbital (120 mg/kg), mice were decapitated. Their spinal cords were dissected in room temperature recording ACSF (in mM: NaCl, 127; KCl, 3; NaH2PO4, 1.2; MgCl2, 1; CaCl2, 2; NaHCO3, 26; D-glucose, 10), with ventral and dorsal roots dissected as distally as possible. To facilitate optogenetic stimulation of dI3 neurons in dI3^CatCh^ mice, dorsal hemisections of lumbar spinal cord segments were made. The ventral hemicord was then allowed to equilibrate in room temperature recording ACSF for at least one hour. Spinal cords were then pinned with the ventral side down to a base of clear Sylgard (Dow Corning) in a recording chamber and perfused with circulating room temperature recording ACSF. Ventral roots were placed in bipolar suction electrodes (A-M Systems Inc.) prior to the optogenetic stimulation protocols. Blue light (470 nm, X-Cite 120LEDmini) was focused on the smallest possible area of the spinal cord using a water-immersed 10x objective lens (Zeiss W N-Achroplan 10x/0.3 M27, 420947-9900), centred on a particular spinal cord segment (identified by the ventral root entry site) between L1 and L5. Each trial stimulated for 10 seconds (continuous light) with a minimum of 10 seconds of control recording after each stimulus.

### EMG recordings

Electrophysiological recordings began 1 week after implanting the EMG electrodes. Recordings were made at the same timepoints as treadmill locomotor assessments. A lightweight cable of 150 cm was attached to the external connector of the EMG implant. This cable fed into a junction box that output to a Model 1700 differential amplifier (A-M systems) and Digidata series 1550B digitizer (Molecular Devices). We measured the motor responses through the implanted EMG electrodes in the tibialis anterior and gastrocnemius muscles of the hindlimbs.

### Perfusion and Immunohistochemistry

Mice were transcardially perfused with ice-cold 4% paraformaldehyde, and their spinal cords were removed and postfixed overnight. The spinal cord was cryoprotected in 30% sucrose, transverse sectioned on a cryostat at (40-50 μm) and collected as free-floating sections. Sections were washed three times with PBS, blocked for one hour in PBS with 0.25% triton-X100 (PBST) and 5% normal goat serum, and then incubated overnight at 4°C with primary antibodies in PBST plus 5% serum. The following day, sections were washed three times with PBS (once with 0.3% H_2_O_2_ to inactivate endogenous peroxidases) and incubated with HRP-conjugated secondary antibodies for three hours at room temperature. Sections were then washed three additional times in PBS prior to incubation for 10 minutes in diluted Opal 690 (Akoya Biosciences). Sections received 2 final PBS washes prior to mounting and coverslipping on Superfrost slides with Immu-Mount. The following antibodies were used for this study: rabbit anti-5-HT2C-R (Immunostar, RRID: AB_572212), rabbit anti-5-HT2A-R (Immunostar, RRID: AB_572211), Goat Anti-Rabbit IgG H&L-HRP (Abcam, ab6721).

### Statistical Analysis

GraphPad Prism 10.4.1 (RRID: SCR_002798) was used to perform statistical analyses and generate the graphs. Data are reported as mean ± S.D and the significance level was set at *p* < 0.05. Comparisons were made using either paired t-tests (two-tailed) or Mixed Model analysis followed by comparison between pairs of groups using Sidak’s multiple comparison test.

## RESULTS

### Acute loss of dI3 excitability results in immediate reduction of locomotor function and motor tone in spinal cord injury mice

We first investigated whether acutely inhibiting dI3 neurons would compromise locomotor function after SCI, as was previously observed when glutamate neurotransmission was developmentally disrupted in dI3 neurons (Bui et al., 2016). Using an intersectional approach to target gene expression to dI3 neurons, we generated tri-hybrid mice that were positive for Isl1-Cre, Vglut2-Flp and CreON-FlpON-hM4Di alleles (Fig 1A). The net effect of this transgenic breeding strategy was the conditional expression of the inhibitory DREADD receptor, hM4Di, in dI3 neurons (which express both Isl1 and Vglut2). These triple transgenic mice are hereafter referred to as dI3^hM4Di^ mice. In dI3^hM4Di^ mice, spinal cord hM4Di expression is localized to dI3 neurons as well as a subset of Vglut2-expressing sensory neurons that are mostly a subset of nociceptors in dorsal root ganglia (Goltash et al., 2025).

**Figure 1.**
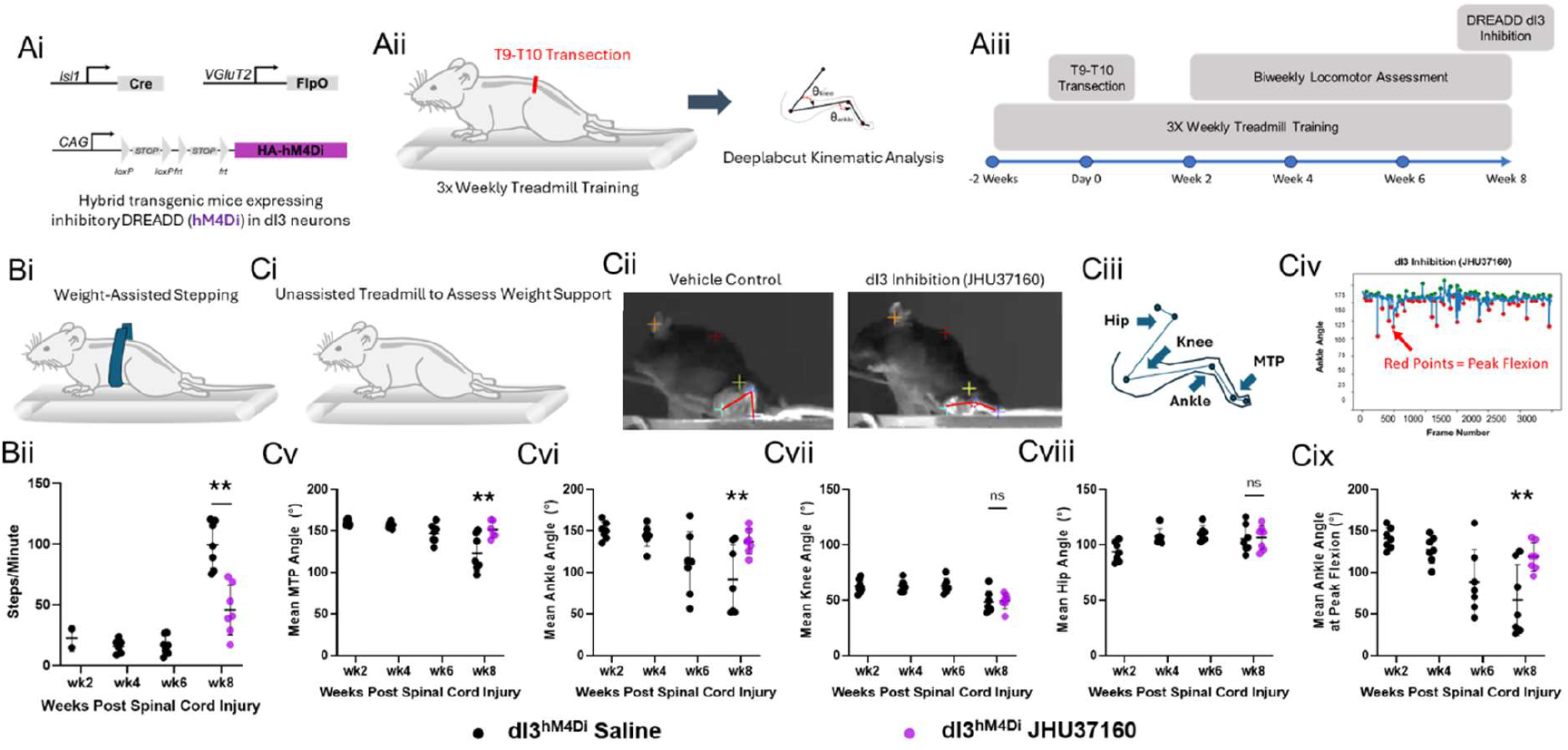
Acute inhibition of dI3 neurons results in the loss of hindlimb stepping and motor tone below the level of injury. Ai) A schematic showing the transgenic expression system for the inhibitory DREADD receptor, hM4Di, in dI3 neurons. Aii) A schematic of the experimental paradigm and Aiii) timeline. Bi) Weight-assistance was provided to spinalized mice to assess stepping on the treadmill. Bii) Treadmill stepping was significantly reduced following dI3 inhibition via the administration of JHU37160 (0.5 mg/kg S.C.) at the experimental endpoint of 8 weeks. Ci-Civ) Unassisted treadmill testing was performed to evaluate the extent of weight support and motor tone in the hindlimbs. Mean and peak joint angles were determined based on the deeplabcut labeling of the iliac crest, hip, knee, ankle, metatarsophalangeal (MTP) joint, and first toe. Cv-Cvi) Distal hindlimb joints, primarily the MTP and ankle joints displayed significant increases in the mean joint angles, corresponding to reduced motor tone and/or weight support through the hindlimbs following the acute inhibition of dI3 neurons. This effect was also observed in relation to the peak ankle angle Cix) and was therefore not merely a result of differences in the number of steps. Cvii-Cviii) Proximal hindlimb joints such as the hip and knee did not show any significant differences in joint angles following dI3 inhibition, possibly due to a lack of meaningful recovery of prior to testing. Statistics were calculated based on paired t-tests (two-tailed) at the 8 week timepoint (n=7). ^*^p<0.05, ^**^p<0.01, ^***^p<0.001, ^****^p<0.0001. Graphs present the mean ± SD.

To prevent any indirect, or chronic effects of dI3 inhibition on the efficacy of locomotor recovery after spinal cord injury, dI3^hM4Di^ mice were not given the DREADD agonist JHU37160 until the final week of the experiment. This allowed the mice time to recover significant locomotor function prior to the dI3 inhibition. Locomotor testing of dI3^hM4Di^ SCI mice shortly after vehicle or JHU37160, a potent DREADD agonist, resulted in the significant reduction of weight-assisted stepping (Figure 1B, paired t-test (two-tailed), n=7, p=0.0024), and the flaccid extension of the distal hindlimb joints of dI3^hM4Di^ SCI mice during unassisted treadmill testing (Figure 1Ci-v, paired t-test (two-tailed), n=7,p_MTP_=0.0034, p_Ankle_=0.0079, p_Knee_=0.8236, p_Hip_=0.8628). In addition to mean joint angles, which largely reflect the resting point of the joint during weight support, the ankle angle at peak flexion was also significantly affected by dI3 inhibition, indicating weaker flexion driven by treadmill-induced sensory stimulation (Figure 1Cvi). While the effects of dI3 inhibition were largely observed in the ankle and metatarsophalangeal joints (Figure 1Cvii-ix), this may have been due to a lack of significant recovery of knee and hip function over the 8 weeks of treadmill training, rather than an indication of specific dI3 neuron action on muscles of the distal hindlimbs.

### dI3 neuron activity is sufficient to drive locomotor activity in ex vivo spinal cord preparations

The above results confirm the prior findings of Bui et al. (2016) pointing to a role of dI3 neurons in the recovery of locomotor function after SCI in mice. However, to date, the primary evidence linking dI3 neurons to locomotor circuits has been derived from loss-of-function experiments (Bui et al., 2013, 2016). It is unclear from these experiments whether the stimulation of dI3 neurons is sufficient to drive locomotor function, or whether their activity merely facilitates or shapes ongoing locomotor activity. We therefore sought to understand the mechanism by which dI3 neurons enable recovery of locomotor function after SCI in mice. First, we tested whether the optogenetic stimulation of dI3 neurons in ex-vivo preparations of the postnatal spinal cord could promote locomotor function. Ventral root recordings of isolated spinal cords of Isl1-Cre; Vglut2-Flp; CreON-FlpON-CatCh mice (herein referred to as dI3^CatCh^ mice) were made after dorsal hemisection to expose dI3 neurons as well as limit optogenetic activation of Isl1^+^/Vglut2^+^ nociceptive afferents (Figure 2A). CatCh is a highly light-sensitive and fast variant of ChR2 (Kleinlogel et al. 2011). Spinal cords were collected from P1-P4 (n=10) dI3^Catch^ mice, and each spinal cord segment between L1 and L5 was exposed to 10-second long, 470 nm illuminations. At each level tested, dI3 illumination resulted in rhythmic, flexor-extensor alternating, locomotor activity in each of the sampled spinal cords (n = 10 out of 10; Figure 2B). The mean frequency of this locomotor activity across all segments was approximately 2.2 Hz (Figure 2B). This locomotor activity induced by optogenetic activation of dI3 neurons suggests that these spinal neurons are capable of driving spinal locomotor networks, at least in neonatal stages.

**Figure 2.**
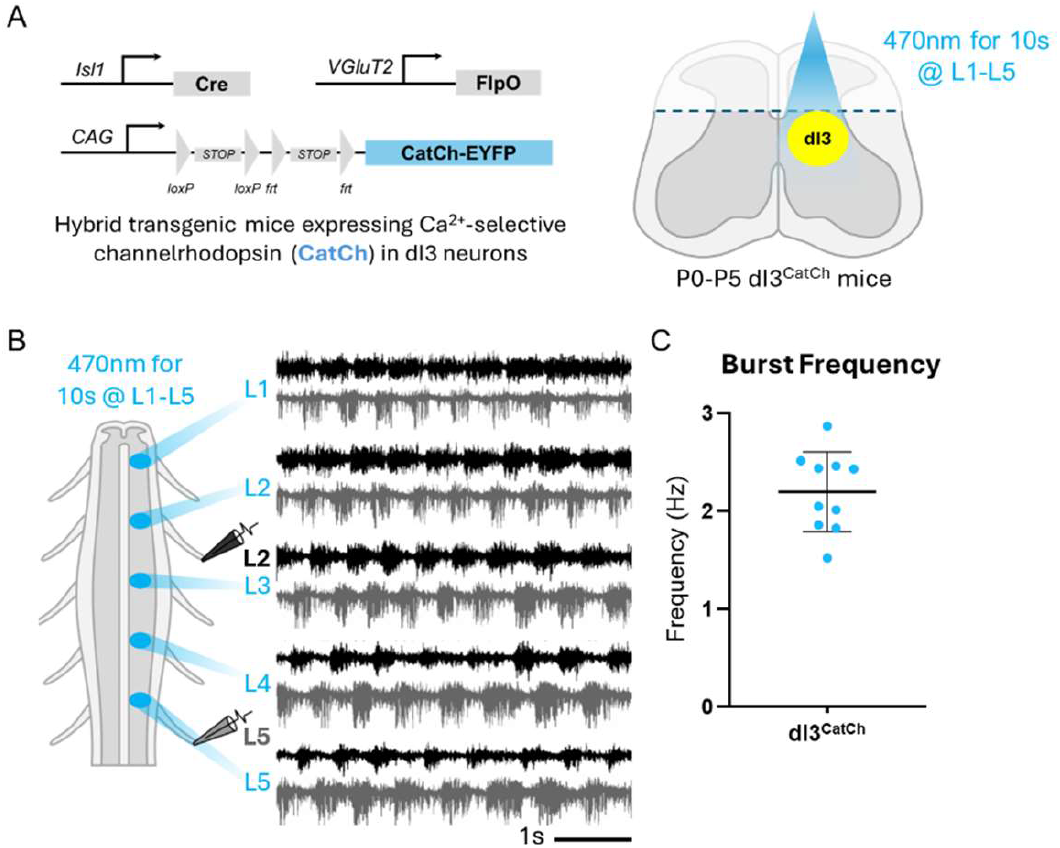
Optogenetic stimulation of dI3 neurons in a neonatal spinal cord preparation generates rhythmic locomotor activity. A) A schematic showing the transgenic expression system for the Calcium selective channelrhodopsin variant, CatCh, in dI3 neurons, and the dorsal horn-removed spinal cord preparation B) Representative recordings from L2 (black) and L5 (grey) ventral roots in response to 10 seconds of continuous 470nm light stimulation. Each row of recordings represents stimulation at a different segment between L1 and L5. C) A plot of the average recorded frequencies of locomotor activity elicited by dI3 neuron stimulation between L1 and L5 (n=10 mice).

### Chemogenetic excitation of dI3 neurons transiently increases hindlimb motor tone during treadmill locomotion and passive stretching

Given that tonic optogenetic stimulation of dI3 neurons was sufficient to drive locomotor activity in neonatal mice, we hypothesized that these neurons provided a similar drive in adult mice. We adopted a less invasive approach to tonically stimulate dI3 neurons and assessed whether chemogenetic excitation of dI3 neurons could similarly elicit locomotor activity in the mice with spinal cord injuries. To test the effects of stimulating dI3 neurons in adult spinalized animals, we expressed the excitatory DREADD receptor, hM3Dq, in spinal Isl1^+^/Vglut2^+^ expressing cells. To prevent activation of Isl1^+^/Vglut2^+^ nociceptors (Goltash et al., 2025), hM3Dq expression in dI3 neurons was achieved by intraspinal injection of an AAV containing a Cre and Flp-dependent hM3Dq construct (Figure 3A). Intraspinal injection of this AAV into Isl1-Cre; Vglut2-Flp animals led to transduction of dI3 neurons (Figure 3A). Two weeks after intraspinal injection of the AAV construct, these mice (referred to hereafter as dI3^hM3Dq^ mice) underwent complete spinal transection at T9-10. A week after the transection, mice were implanted with EMG electrodes in the left and right ankle flexor, tibialis anterior, and ankle extensor, gastrocnemius, for longitudinal EMG recordings (Figure 3A).

**Figure 3.**
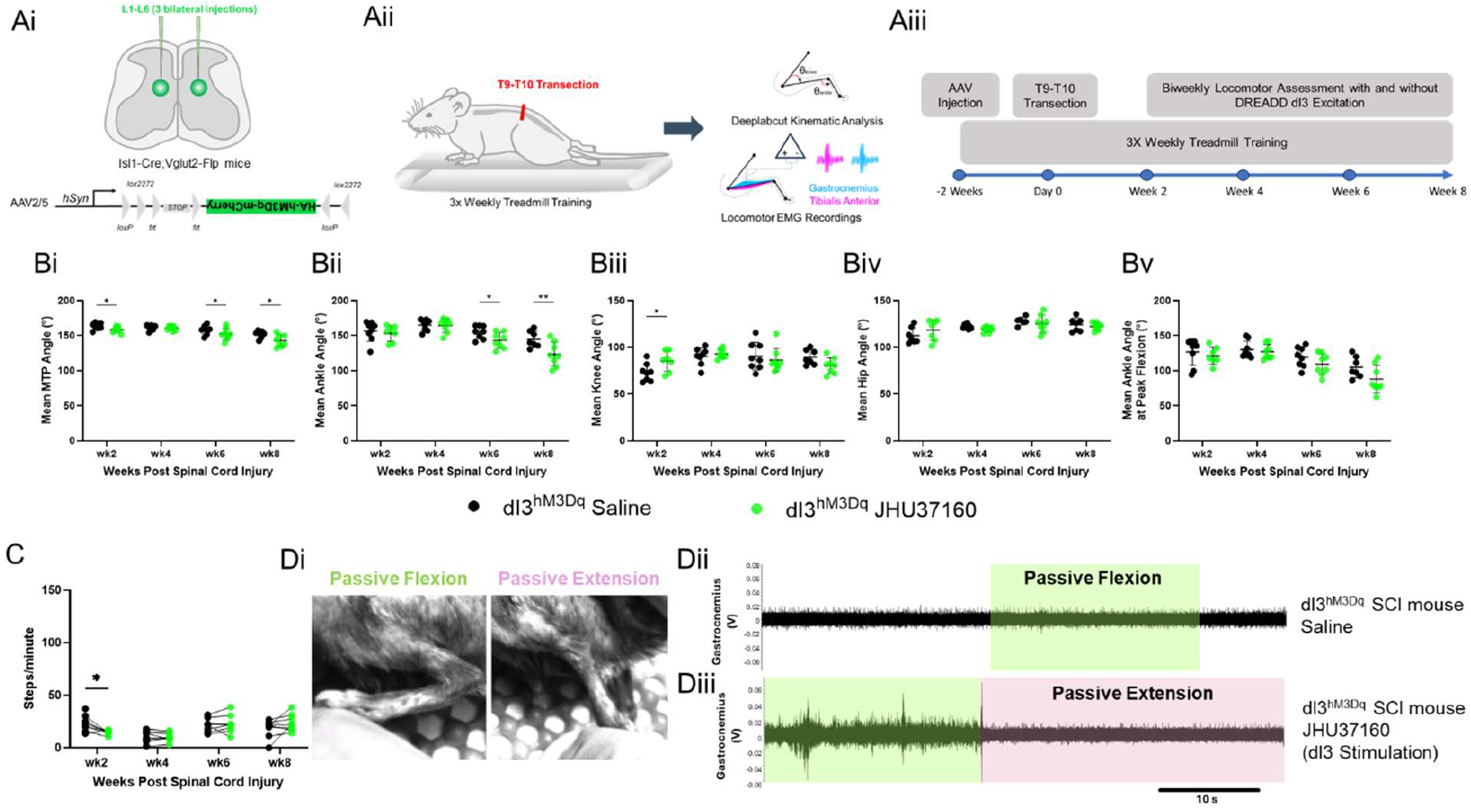
Chemogenetic stimulation of dI3 neurons transiently increases distal hindlimb joint motor tone and resistance to static stretch. Ai) A schematic representing the methodology used to express the excitatory DREADD receptor hM3Dq. An AAV designed to express an HA-tagged, hM3Dq-mCherry fusion protein in the presence of Cre and Flp recombinases was intraspinally injected bilaterally via the three consecutive intralaminar spaces to target spinal segments corresponding to L1-L6. Aii-iii) Mice expressing hM3Dq then received a T9-T10 spinal cord transection, and surgical implantation of EMG electrodes into the gastrocnemius and tibialis anterior muscles. Mice received three weekly treadmill training sessions and biweekly assessment of locomotor function in the presence and absence of the DREADD agonist, JHU37160 (0.5 mg/kg S.C.). B) Mean joint angles were assessed across both conditions. Transient increases in joint flexion were observed at multiple timepoints for the Bi) MTP and Bii) ankle following the chemogenetic stimulation of lumbar dI3 neurons. Biii) Knee and Biv) hip angles were largely unaffected by the stimulation of dI3 neurons, except for a slight increase in knee extension at the week 2 timepoint. C) Treadmill stepping was modestly decreased at week 2, but was otherwise unaffected by dI3 stimulation. Di) Passive stretching of the mouse hindlimb was performed in conjunction with EMG recordings to assess motor tone before and after dI3 stimulation. Dii-iii) EMG activity in the gastrocnemius in response to manual ankle flexion was significantly higher following chemogenetic stimulation of dI3 neurons than in the saline control condition. Statistics were calculated based on Linear Mixed Model analysis with a Sidak’s multiple comparisons test (n=8). ^*^p<0.05, ^**^p<0.01, ^***^p<0.001, ^****^p<0.0001. Graphs present the mean ± SD.

Biweekly unassisted treadmill locomotor assessments revealed that the administration of the DREADD agonist JHU37160 (0.5 mg/kg, S.C.) to stimulate dI3s resulted in relatively modest increases in mean flexor tone at the MTP and ankle joints, particularly at weeks 6 and 8 post-injury (Figure 3Bi-ii, Mixed Model Analysis, Sidak’s test, n=8, p_MTP-6wk_=0.0130, p_MTP-8wk_=0.0259, p_Ankle-6wk_=0.0278, p_Ankle-8wk_=0.0098). These effects were not observed at the knee and hip joints (Figure 3Biii-iv). Ankle angle at peak flexion did appear to be modified by JHU37160 administration (Figure 3Bv, Mixed Model Analysis, n=8, p_treatment-effect_=0.0383), but no specific timepoint reached the threshold for statistical significance. However, contrary to expectation, there was no significant increase in weight-supported stepping observed after dI3 stimulation (Figure 3C). Indeed, the only significant difference was an apparent decrease in stepping after the administration of the first JHU37160 dose at week 2 (Figure 3C, Mixed Model Analysis, Sidak’s test, n=8, p_Steps-2wks_=0.0233).

Given that the main effect of lumbar dI3 neuron stimulation was related to distal joint angles during unassisted treadmill testing, we hypothesized that perhaps dI3 activity was enhancing the motor response to passive stretch induced by the movement of the treadmill. To test this hypothesis, we subjected the hindlimbs of SCI dI3^hM3Dq^ animals to manual limb extension or flexion (Figure 3Di). When vehicle control was administered to SCI dI3^hM3Dq^ mice, manual limb extension or flexion had no effect on EMG activity (Figure 3Dii). In contrast, administration of JHU37160 led to EMG responses consistent with the activation of stretch responses in either gastrocnemius when the ankle was flexed, or tibialis anterior when the ankle was extended (Figure 3Diii, supplementary video 1).

### Chronic effects of dI3 stimulation limit natural locomotor recovery and weight support

In addition to the transient effects of chemogenetic dI3 stimulation described above, some persistent effects on the locomotor phenotype were also noted. Primarily, it was observed that the dI3 stimulation cohort did not experience the same recovery trajectory in relation to both weight-assisted stepping and hindlimb posture in the unassisted treadmill testing. When comparing the saline injected trials between dI3^hM4Di^ mice and dI3^hM3Dq^ mice, dI3^hM3Dq^ mice that had previously received chemogenetic dI3 stimulation performed significantly worse at weight-assisted stepping at 8 weeks post-injury (Figure 4A, Mixed Model Analysis, Sidak test, n_hM4Di_=7, n_AAV-hM3Dq_=8, p_Wk8_<0.0001). These mice also demonstrated increased mean MTP (Figure 4Bi, Mixed Model Analysis, Sidak test, n_hM4Di_=7, n_AAV-hM3Dq_=8, p_Wk8_<0.0001) and ankle angles (Figure 4Bii, Mixed Model Analysis, Sidak test, n_hM4Di_=7, n_AAV-hM3Dq_=8, p_Wk6_=0.0011, p_Wk8_<0.0001). Peak ankle flexion was also notably reduced in the dI3^hM3Dq^ mice (Figure 4Biii, Mixed Model Analysis, Sidak test, n_hM4Di_=7, n_AAV-hM3Dq_=8, p_Wk8_<0.05) relative to the dI3^hM4Di^ mice. In aggregate, these measures indicate that semi-regular chemogenetic stimulation of dI3 neurons paradoxically disrupts the typical locomotor recovery observed after spinal cord injury.

**Figure 4.**
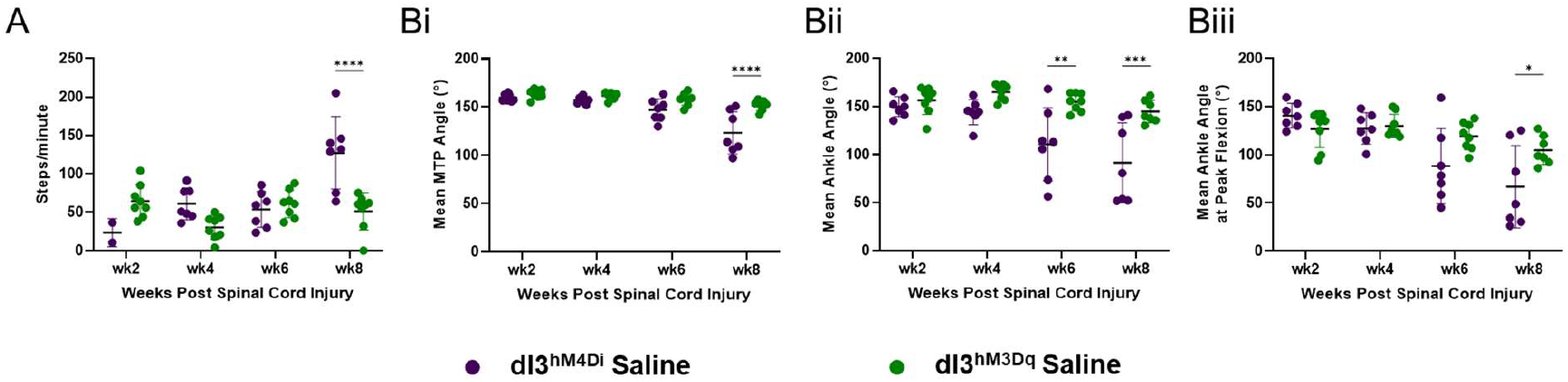
Regular dI3 stimulation using DREADD receptor hM3Dq results in chronic behavioural changes that limit locomotor functional recovery after spinal cord injury. A) Weight supported stepping was reduced in mice that had received regular dI3 chemogenetic stimulation by week 8 post-injury. Bi-ii) Mean MTP and ankle angles were also more extended during unassisted treadmill testing in the group that had previously received dI3 stimulation by 8 weeks post-injury. Biii) Peak ankle angle was similarly affected, demonstrating that ankle flexion during stepping was reduced as well in dI3 stimulation group by 8 weeks post-injury. Statistics were calculated based on Linear Mixed Model analysis with a Sidak’s multiple comparisons test (n_hM4Di_=7, n_AAV-hM3Dq_=8). ^*^p<0.05, ^**^p<0.01, ^***^p<0.001, ^****^p<0.0001. Graphs present the mean ± SD.

### dI3 expression of 5-HT2C receptor provides potential link to homeostatic mechanisms governing dI3 activity after spinal cord injury

Given that acute dI3 inhibition and chronic dI3 stimulation produced similar effects on locomotion following spinal cord injury, we hypothesized that these paradoxical findings may be the result of homeostatic mechanisms governing the excitability of neurons. As it is known that the 5-HT2C receptor has a constitutively active isoform that is expressed after spinal cord injury (Fouad et al., 2010; Murray et al., 2010, 2011; Ren et al., 2013), we examined 5-HT2C receptor expression in the spinal cords of intact and spinal cord injury mice. Critically, we found that the majority of dI3 neurons (Figure 5A-C, 49/50 intact, 201/219 SCI) robustly expressed the 5-HT2C-receptor. Further, we found that in samples having received chemogenetic dI3 stimulation, the dI3 localization of 5-HT2C expression was membrane-bound (Figure 5Bi-iv) – consistent with the pre-injury expression pattern – and in contrast to the clumped, internalized expression pattern observed in the constitutively active isoform (Murray et al., 2010). As the expression of the constitutively active 5-HT2C receptor isoform would increase dI3 excitability, its presence in unstimulated samples, and absence in stimulated samples, could provide a basis for the muted locomotor recovery in dI3^hM3Dq^ mice after spinal cord injury.

**Figure 5.**
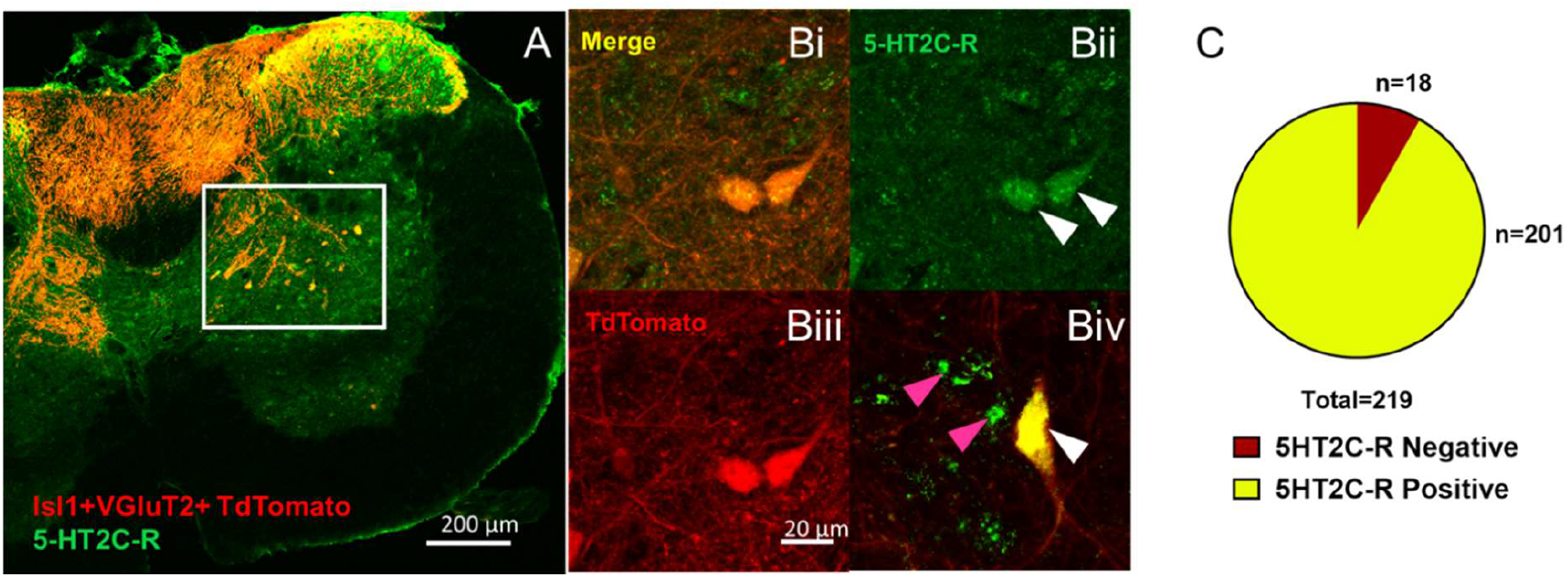
dI3 neurons exhibit normal membrane-bound 5HT2C receptor phenotype when chemogenetic stimulation is applied after spinal cord injury. A and C) The majority of dI3 neurons strongly express the 5-HT2C receptor following spinal cord injury. Bi-iv) dI3 neurons (tdTomato+) from SCI mice that received chemogenetic stimulation exhibit the normal membrane-bound pattern of expression (white arrows). Clumped, internalized expression of the 5-HT2C receptor has been linked to the constitutively-active isoform that emerges post-spinal cord injury (pink arrows).

### Combination of chemogenetic dI3 stimulation and quipazine administration provide an additive benefit on treadmill locomotion

While dI3 stimulation in neonatal spinal cords was a potent driver of locomotor function, chemogenetic stimulation of dI3 neurons alone did not lead to statistically significant increases in stepping function when compared to vehicle-treated SCI dI3^hM3Dq^ animals. We therefore tested whether concurrent administration of the serotonergic agonist quipazine and hM3Dq activation could lead to greater stepping function. Quipazine has been previously used as a neuromodulator for increased stepping function after SCI in animal models. At the selected dose (0.5 mg/kg), it has been reported that quipazine facilitates locomotor activity without directly initiating locomotor rhythm, allowing the locomotor networks to respond to sensory cues provided by the treadmill (Fong et al., 2005). We observed that concurrent administration of quipazine and increased dI3 IN activity by JHU37160 administration in SCI dI3^hM3Dq^ animals improved weight-assisted stepping function over vehicle or quipazine-alone treated SCI dI3^hM3Dq^ animals at all timepoints (2, 4, 6, and 8 weeks) post spinal cord injury (Figure 6B, Mixed Model Analysis, Sidak’s test, n=7, p_Wk2_=0.0484, p_Wk4_=0.0499, p_Wk6_=0.0320, p_Wk8_=0.0105). While both the combined treatment, and quipazine alone produced statistically significant changes to the mean joint angles for MTP (Figure 6Ci), ankle (Figure 6Cii) and knee (Figure 6Ciii), there was no clear distinction between the two treatment paradigms. This was also the case for the mean ankle angle at peak flexion (Figure 6Cv).

**Figure 6.**
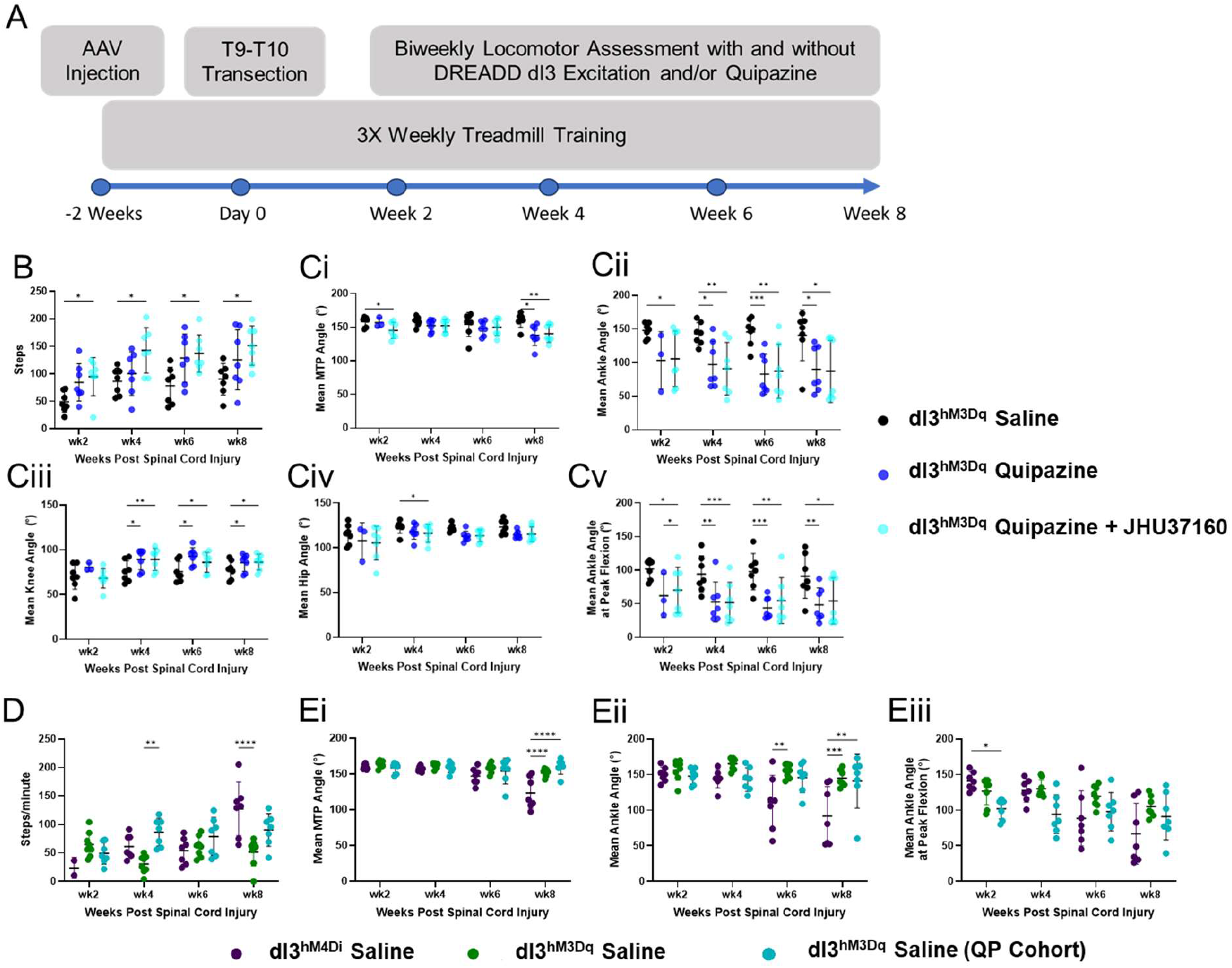
Combination of type 2 serotonin receptor agonist, quipazine, with dI3 chemogenetic stimulation provides a combinatorial benefit on locomotor function after SCI, and reduces chronic effects of dI3 stimulation on locomotor stepping. A) A schematic of the experimental timeline. B) Weight assisted treadmill stepping was significantly increased in mice receiving a combination of quipazine (0.5 mg/kg I.P.) and JHU37160 (0.5 mg/kg S.C.) relative saline control injections. This was found across all timepoints measured. Ci-Cv) Mean joint angles for most of the hindlimb joints measured during unassisted treadmill walking were significantly impacted by the administration of quipazine and quipazine + JHU37160. However, few timepoints demonstrated evidence of an additive effect of quipazine + JHU37160 (Ci – MTP week 2, Cii – ankle week 2, Civ – Hip week 4). D) Chronic effects of dI3 stimulation on weight-assisted stepping were mitigated by the addition of quipazine to the treatment paradigm. Ei-Eiii) Chronic effects of dI3 stimulation on mean joint angles were not rescued by the addition of quipazine. Statistics were calculated based on Linear Mixed Model analysis with a Sidak’s multiple comparisons test (n_hM4Di_=7, n_AAV-hM3Dq_=8, n_AAV-hM3Dq (QP Cohort)_=7). ^*^p<0.05, ^**^p<0.01, ^***^p<0.001, ^****^p<0.0001. Graphs present the mean ± SD.

In terms of the chronic effects of the combined quipazine and chemogenetic dI3 stimulation treatment strategy, the deficits in weight-assisted stepping associated with dI3 stimulation alone were substantially rescued by the addition of quipazine, as the saline condition of the dI3^hM4Di^ cohort was not significantly different from saline condition of the dI3^hM3Dq^ + quipazine cohort (Figure 6D, Mixed Model Analysis, Sidak’s test, n=7, p_Wk8_=0.1090). However, these benefits were limited to weight-assisted stepping, as the mean joint angles were not significantly rescued during unassisted treadmill locomotion (Figure 6E).

The sum of our experiments suggests that dI3 neurons can increase activation of spinal locomotor circuits either by direct activation of spinal locomotor circuits or indirect activation of spinal locomotor circuits through increased proprioceptive sensorimotor function at the level of the spinal cord in spinalized animals.

## DISCUSSION

Unlike primates that show limited capacity for recovery of lost motor function after spinal cord injury, many mammalian species can exhibit locomotor activity in certain settings. Given the proper sensory, chemical, or electrical stimulation, these species can show the hallmarks of locomotor activity such as rhythmic stepping with proper left-right and swing-stance alternations, even following complete transection of the spinal cord. The mechanisms by which the spinal cord can produce locomotor activity without descending supraspinal commands are gradually being revealed. For instance, sensory information from muscle and cutaneous afferents have been shown to be critical for recovery of proper locomotor activity (Bouyer and Rossignol, 2003; Sławińska et al., 2012; Takeoka et al., 2014; Takeoka and Arber, 2019). A recent study using RNA sequencing of the spinal cord after spinal cord injury has identified V2a interneurons as being of importance to locomotor activity after spinal cord injury (Kathe et al., 2022). These predominantly ipsilateral excitatory spinal interneurons are shown to be critical for rhythmogenesis in zebrafish (Eklöf-Ljunggren et al., 2012; Ampatzis et al., 2014; Ljunggren et al., 2014) and ensure proper left-right coordination in the mouse spinal cord at certain speeds (Crone et al., 2008).

dI3 neurons were identified as candidates involved in locomotor recovery due to their integration of proprioceptive and mechanoreceptive inputs, as well as their glutamatergic connections to spinal motoneurons and potentially to spinal locomotor circuits (Bui et al., 2013, 2016). Developmentally regulated silencing of their glutamatergic neurotransmission – achieved through Isl1-driven Cre-mediated deletion of Vglut2 – indicated that these neurons are important for the recovery of locomotor activity following spinalization (Bui et al., 2016). In this study, we used a combination of optogenetic and chemogenetic approaches to provide further evidence that these spinal neurons can directly excite spinal locomotor circuits. While the effect on locomotor output was substantially stronger using optogenetic stimulation in neonatal preparations than in the *in vivo* chemogenetic stimulation protocol, possibly due to the greater synchronicity and strength of stimulation, the results support that dI3 neurons could form part of the sensorimotor circuitry that activates locomotor circuits after spinal cord injury.

However, the dominant effect of dI3 modulation after spinal cord injury appeared to be related to the regulation of motor tone. Definitionally, motor tone relates to the inherent involuntary tension that a muscle actively maintains at rest. It provides resistance to passive stretch and is critical for the maintenance of posture and reflexes. In the context of this work, motor tone was largely assessed by examining the postural changes on a treadmill task following dI3 modulation, or via passive manipulation of the hindlimbs during EMG recordings. In both cases, we observed results consistent with the hypothesis that dI3 activity promotes hindlimb motor tone after spinal cord injury, namely, increased hindlimb flaccidity after dI3 inhibition, and increased responsiveness to passive stretch in the primary ankle flexor and extensor muscles. The main complication to this interpretation is the paradoxical reduction of motor tone in mice with prior dI3 stimulation. While the exact mechanism responsible for this effect has yet to be determined, it could be speculated that this result could be the consequence of either excessive stimulation, or intracellular signaling of the hM3Dq GPCR, interfering with the homeostatic mechanisms responsible for increasing dI3 excitability after SCI.

The relationship between motor tone and locomotor recovery after spinal cord injury is intriguing, in that the dysregulation of motor tone is often a significant hurdle for spinal cord injury patients. Low tone, or hypotonia, results in flaccidity, poor weight support, and the attenuation of reflexes. It is generally a symptom of the early stages of spinal cord injury, and is frequently replaced by hypertonia – manifesting as spasticity (velocity-dependent) or rigidity (velocity-independent). While hypertonia poses its own problems for functional recovery, lack of tone largely prevents motor recovery, as flaccid muscles offer relatively little proprioceptive feedback, a critical component of physical rehabilitation (Capogrosso et al., 2013; Takeoka et al., 2014; Takeoka and Arber, 2019). Indeed, it has been hypothesized previously that hypertonia after spinal cord injury is perhaps a semi-beneficial homeostatic response to a lack of sufficient input to spinal cord circuits (reviewed in D’Amico et al., 2014). This point of view has circumstantial support from reports of impacts on locomotor recovery from anti-spastic interventions (Murray et al., 2010, 2011; Angeli et al., 2012).

One well-described link between locomotor recovery and hypertonia/spasticity after spinal cord injury involves the 5-HT2C receptor. Seminal works from Murray et al (2010,Murray et al 2011) and Fouad et al (2010), collectively discovered that a constitutively active form of the 5-HT2C receptor was not only responsible for rescuing the persistent inward currents necessary for robust motoneuron activity needed to sustain locomotor activity, but was also largely responsible for the uncontrolled involuntary spasms present in spastic animals. While these studies examined the 5-HT2C receptor in the context of spinal motoneurons, it should be noted that motoneurons have been reported to express 5-HT2C receptors relatively late, potentially months after spinal cord injury in rats (Murray et al., 2010; Ren et al., 2013). This, of course, extends far past the timeline when locomotor recovery begins in rats, with gradual improvements in motor tone and movement expected between 2-6 weeks post-injury (Alluin et al., 2011). As such, other neuron populations, such as the dI3 neurons which express the 5-HT2C receptor, may be responsible for the earlier effects on motor tone, posture, and locomotor improvement.

In summary, our results in neonatal mice suggest that dI3 neurons are capable of activating spinal locomotor circuits. However, while inhibiting these neurons impacts negatively the recovery of locomotor function after spinal cord injury, the chronic stimulation of dI3 neurons does not have the converse beneficial effects that we hypothesized. The expression of different forms of 5-HT2C receptors on dI3 neurons may be indicative of complex homeostatic regulation of dI3 neuron activity after spinal cord injury that could tilt the balance between hypotonia versus hypertonia. Future studies evaluating how to modulate this homeostatic regulation could be invaluable in guiding therapeutic approaches to promote recovery of locomotor function through dI3 neuron modulation.

## REFERENCES

Akay T, Acharya HJ, Fouad K, Pearson KG (2006) Behavioral and Electromyographic Characterization of Mice Lacking EphA4 Receptors. J Neurophysiol 96:642–651.

Alluin O, Karimi-Abdolrezaee S, Delivet-Mongrain H, Leblond H, Fehlings MG, Rossignol S (2011) Kinematic Study of Locomotor Recovery after Spinal Cord Clip Compression Injury in Rats. J Neurotrauma 28:1963–1981.

Ampatzis K, Song J, Ausborn J, El Manira A (2014) Separate Microcircuit Modules of Distinct V2a Interneurons and Motoneurons Control the Speed of Locomotion. Neuron 83:934–943.

Angeli C, Ochsner J, Harkema S (2012) Effects of chronic baclofen use on active movement in an individual with a spinal cord injury. Spinal Cord 50:925–927.

Barbeau H, Rossignol S (1987) Recovery of locomotion after chronic spinalization in the adult cat. Brain Res 412:84–95.

Bouyer LJ, Rossignol S (2003) Contribution of cutaneous inputs from the hindpaw to the control of locomotion. II. Spinal cats. J Neurophysiol 90:3640–3653.

Bui TV, Akay T, Loubani O, Hnasko TS, Jessell TM, Brownstone RM (2013) Circuits for grasping: spinal dI3 interneurons mediate cutaneous control of motor behavior. Neuron 78:191–204.

Bui TV, Stifani N, Akay T, Brownstone RM (2016) Spinal microcircuits comprising dI3 interneurons are necessary for motor functional recovery following spinal cord transection. eLife 5:1–20.

Capogrosso M, Wenger N, Raspopovic S, Musienko P, Beauparlant J, Bassi Luciani L, Courtine G, Micera S (2013) A Computational Model for Epidural Electrical Stimulation of Spinal Sensorimotor Circuits. J Neurosci 33:19326–19340.

Crone SA, Quinlan KA, Zagoraiou L, Droho S, Restrepo CE, Lundfald L, Endo T, Setlak J, Jessell TM, Kiehn O, Sharma K (2008) Genetic Ablation of V2a Ipsilateral Interneurons Disrupts Left-Right Locomotor Coordination in Mammalian Spinal Cord. Neuron 60:70–83.

D’Amico JM, Condliffe EG, Martins KJB, Bennett DJ, Gorassini MA (2014) Recovery of neuronal and network excitability after spinal cord injury and implications for spasticity. Front Integr Neurosci 8 Available at: https://www.frontiersin.org/journals/integrative-neuroscience/articles/10.3389/fnint.2014.00036/full [Accessed September 2, 2025].

Edgerton VR, Roy RR (2009) Activity-dependent plasticity of spinal locomotion: implications for sensory processing. Exerc Sport Sci Rev 37:171–178.

Eklöf-Ljunggren E, Haupt S, Ausborn J, Dehnisch I, Uhlén P, Higashijima S, El Manira A (2012) Origin of excitation underlying locomotion in the spinal circuit of zebrafish. Proc Natl Acad Sci U S A 109:5511–5516.

Fong AJ, Cai LL, Otoshi CK, Reinkensmeyer DJ, Burdick JW, Roy RR, Edgerton VR (2005) Spinal Cord-Transected Mice Learn to Step in Response to Quipazine Treatment and Robotic Training. J Neurosci 25:11738–11747.

Fouad K, Rank MM, Vavrek R, Murray KC, Sanelli L, Bennett DJ (2010) Locomotion after spinal cord injury depends on constitutive activity in serotonin receptors. J Neurophysiol 104:2975–2984.

Goltash S, Khodr R, Bui TV, Laliberte AM (2025) An optogenetic mouse model of hindlimb spasticity after spinal cord injury. Exp Neurol 386:115157.

Kathe C et al. (2022) The neurons that restore walking after paralysis. Nature 611:540–547.

Lagerstrom MC, Rogoz K, Abrahamsen B, Persson E, Reinius B, Nordenankar K, Olund C, Smith C, Mendez JA, Chen ZF, Wood JN, Wallen-Mackenzie A, Kullander K (2010) VGLUT2-dependent sensory neurons in the TRPV1 population regulate pain and itch. Neuron 68:529–542.

Ljunggren EE, Haupt S, Ausborn J, Ampatzis K, El Manira A (2014) Optogenetic activation of excitatory premotor interneurons is sufficient to generate coordinated locomotor activity in larval zebrafish. J Neurosci Off J Soc Neurosci 34:134–139.

Mathis A, Mamidanna P, Cury KM, Abe T, Murthy VN, Mathis MW, Bethge M (2018) DeepLabCut: markerless pose estimation of user-defined body parts with deep learning. Nat Neurosci 21:1281–1289.

Murray KC, Nakae A, Stephens MJ, Rank M, D’Amico J, Harvey PJ, Li X, Harris RLW, Ballou EW, Anelli R, Heckman CJ, Mashimo T, Vavrek R, Sanelli L, Gorassini MA, Bennett DJ, Fouad K (2010) Recovery of motoneuron and locomotor function after spinal cord injury depends on constitutive activity in 5-HT2C receptors. Nat Med 16:694–700.

Murray KC, Stephens MJ, Ballou EW, Heckman CJ, Bennett DJ (2011) Motoneuron Excitability and Muscle Spasms Are Regulated by 5-HT2B and 5-HT2C Receptor Activity. J Neurophysiol 105:731–748.

Nath T, Mathis A, Chen AC, Patel A, Bethge M, Mathis MW (2019) Using DeepLabCut for 3D markerless pose estimation across species and behaviors. Nat Protoc 14:2152–2176.

Ren L-Q, Wienecke J, Chen M, Møller M, Hultborn H, Zhang M (2013) The time course of serotonin 2C receptor expression after spinal transection of rats: An immunohistochemical study. Neuroscience 236:31–46.

Santos ACR dos, Laurindo RP, Pestana FM, Heringer L dos S, Canedo NHS, Martinez AMB, Marques SA (2024) Exercise Volume Can Modulate the Regenerative Response to Spinal Cord Injury in Mice. Neurotrauma Rep 5:721–737.

Sławińska U, Majczyński H, Dai Y, Jordan LM (2012) The upright posture improves plantar stepping and alters responses to serotonergic drugs in spinal rats. J Physiol 590:1721–1736.

Takeoka A, Arber S (2019) Functional Local Proprioceptive Feedback Circuits Initiate and Maintain Locomotor Recovery after Spinal Cord Injury. Cell Rep 27:71-85.e3.

Takeoka A, Vollenweider I, Courtine G, Arber S (2014) Muscle spindle feedback directs locomotor recovery and circuit reorganization after spinal cord injury. Cell 159:1626–1639.

